# Tendon-derived injectable bio-instructive gel augments tenogenic differentiation of iMSC-SCX cells and extracellular matrix remodeling

**DOI:** 10.64898/2026.09.01.748431

**Authors:** Ahmet Pazarceviren, Hyeongseop Keum, Léo-Paul Tricou, Minoo Bastani, Melissa Chavez, Julia Sheyn, Gaia Dupont, Myung-Seo Kim, Wafa Tawackoli, Benjamin Freedman, Dmitriy Sheyn

## Abstract

In the US, 33 million musculoskeletal injuries have been reported per year, with 50% involving tendons and ligaments in both athletic and aging populations. Tendon repair often results in the formation of biomechanically inferior scar tissue rather than functional regeneration. Local cell therapy is garnering significant interest in tendon repair because it provides a targeted means to repopulate defects with potent therapeutic cells. Here, we developed a porcine tendon-derived thermoresponsive extracellular matrix hydrogel (TG) as an injectable, bio-instructive carrier for scleraxis-overexpressing induced mesenchymal stem cell–derived tenocytes (iTenocytes), with the goal of improving cell retention and overall transplantation success. TG was compositionally distinct from purified collagen (PC, from rat tail type 1 collagen) and exhibited favorable material properties for a minimally invasive percutaneous strategy, including thermoresponsive gelation, shear-thinning injectability, retention at the injection site, and controlled biodegradation. *In vitro*, TG supported three-dimensional cell residence, promoted cell interconnectivity and redistribution at the matrix interface, and increased collagen type I release from iTenocytes compared to those embedded in PC. Transcriptomic and proteomic analyses further showed that TG enhanced programs associated with tenogenic maturation, extracellular matrix assembly, focal adhesion, mechano-transduction, and remodeling. In a rat Achilles tendon partial defect model, both optical imaging and histological analyses demonstrated retention of iTenocytes at the injection site. Additionally, confocal imaging demonstrated retention of iTenocytes within the defect site for up to 10 days. Together, these findings identify TG as a biofunctional injectable carrier that supports the tenogenic characteristics of iTenocytes and enhances their local persistence after transplantation.

## 1. Introduction

Tendon injuries, ranging from acute ruptures to chronic tendinopathies, represent a major clinical and socioeconomic burden. Tendon-related pathologies account for a substantial proportion of musculoskeletal injuries, with increasing prevalence linked to aging and high-impact physical activity [1]. Despite conservative care and surgical repair strategies, including physical therapy, autografts, and synthetic sutures, clinical outcomes remain limited by incomplete restoration of native tendon structure and mechanics, high reinjury risk, and fibrotic scar formation [2–4].

A major barrier to functional tendon regeneration is the native tendon environment, which is hypovascular and sparsely cellular [5]. These features restrict oxygen, nutrient, and progenitor-cell supply to the injury site, contributing to poor intrinsic healing. Pharmacological approaches and injectable biologics such as platelet-rich plasma (PRP) have shown inconsistent efficacy, in part because delivered factors degrade rapidly and lack a stable matrix for sustained local activity [6, 7]. These limitations have motivated cell-based strategies aimed at replenishing the cellular pool and modulating the injury microenvironment toward regeneration.

Conventional stem cell sources for tendon repair, including bone marrow–derived mesenchymal stem cells (BM-MSCs) and adipose-derived stem cells, are limited by phenotypic heterogeneity, donor variability, cellular senescence, and inconsistent tenogenic potential [8–10]. Additionally, off-target differentiation toward bone or cartilage lineages may further compromise tendon mechanics [9]. To address these limitations, we recently developed scleraxis-overexpressing human induced mesenchymal stem cells (iMSC-SCX), referred to here as iTenocytes. Scleraxis (SCX), a basic helix–loop–helix transcription factor and key regulator of tenogenesis, promotes tendon-lineage programming in iPSC-derived MSCs. Compared with primary BM-MSCs, iTenocytes show enhanced tenogenic potential, including increased expression of late tendon markers such as tenomodulin (TNMD) and Mohawk (MKX) [11]. Nevertheless, effective therapeutic use of these cells remains constrained by poor retention and potential viability loss after delivery [12–15].

Biomaterial carriers can improve local cell delivery, but many existing systems lack tendon-specific instructive cues. Synthetic polymers such as PEG or PLGA provide tunable delivery platforms but generally require exogenous bioactive supplementation to support tendon-lineage maintenance [16]. Natural hydrogels such as collagen type I and fibrin offer improved biocompatibility, yet they do not fully recapitulate the biochemical complexity of native tendon extracellular matrix (ECM) [17]. Thus, there remains a need for an injectable carrier that protects cells during delivery, promotes local retention, and provides tendon-relevant cues that synergize with SCX-driven tenogenic programming.

To meet this need, we developed a porcine tendon-derived decellularized ECM hydrogel, referred to as TG, as an injectable, bio-instructive cell carrier. TG is generated by decellularization and solubilization of tendon tissue, preserving tendon-associated ECM components, including collagens, fibronectin, tenascin-C, and signaling molecules such as TGFβ isoforms [18–23]. In addition to its biochemical composition, TG exhibits clinically useful thermoresponsive behavior: it remains liquid at room temperature, enabling cell encapsulation with minimal handling stress, and undergoes spontaneous gelation at 37°C to form a stable fibrous matrix [24, 25]. This sol-to-gel transition allows TG to conform to irregular tendon defects, support localized matrix formation, and reduce cell leakage after implantation.

In this study, we integrated iTenocytes with TG to create an injectable, bio-instructive cell delivery platform. We tested whether TG could maintain tenogenic features of iTenocytes in vitro and improve their local retention after transplantation into a rat Achilles tendon partial defect. Together, these studies evaluate TG as a biofunctional matrix for iTenocyte delivery.

## 2. Results

### 2.1. TG retains tendon-associated ECM composition after decellularization

To develop an injectable and bio-instructive cell carrier system, TG was generated from porcine Achilles tendon **(Fig. 1a)**. Successful decellularization of tendon tissues was assessed using hematoxylin and eosin (H&E) staining, which revealed a marked reduction in cellular content in the TG post-decellularization compared with the fresh tendon **(Fig. 1b)**. Consistently, DNA quantification demonstrated substantially lower DNA levels in the TG group relative to the fresh tendon (**Fig. 1C**). Proteomic profiling of TG was performed in comparison to commercially available purified collagen (PC; from rat tail type 1 collagen), showing distinct protein abundance patterns between the two groups (**Fig. S1**). Analysis of collagen composition showed that both materials were predominantly composed of collagen type 1 (COL1), along with contributions from other collagen subtypes and ECM-associated proteins (**Fig. 1d**). Notably, grouped analysis of tendon-healing associated collagens [20], including COL1A1, COL1A2, and COL3A1, demonstrated that TG exhibited a higher COL1-to-COL3 ratio within their groups (**Fig S2**). The relative collagen composition differed significantly between TG and PC (χ²(3) = 11.94, p = 0.0076). TG exhibited a higher relative contribution of COL1A2 and lower contributions of COL3A1 and other collagen-associated proteins compared with PC. Functional enrichment analysis of differentially represented proteins identified pathways such as ECM– receptor interaction, protein digestion and absorption, cytoskeletal organization in muscle cells, TGF-β signaling, and focal adhesion (**Fig. 1e**).

**Figure 1.**
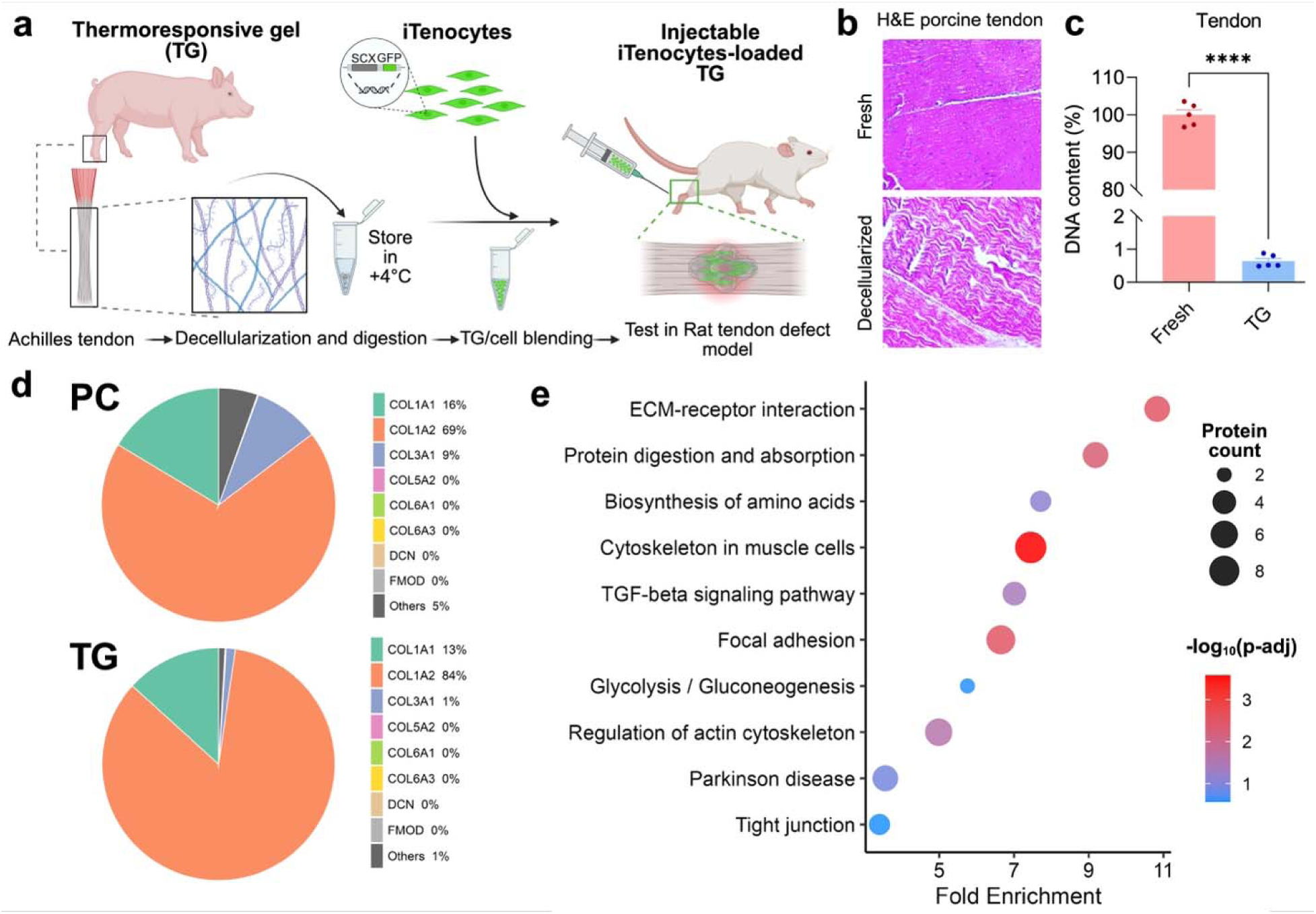
Porcine Achilles tendon ECM-derived thermoresponsive gel (TG) is decellularized and compositionally distinct from commercially available purified collagen (PC). a) Schematic workflow showing porcine Achilles tendon harvesting, decellularization and digestion to generate TG, storage at 4 °C, mixing with iTenocytes, and injection of iTenocyte-loaded TG for testing in a rat tendon defect model. b) Representative H&E images of fresh tendon and TG sections. Scale bars, 300 µm. Data are presented as mean ± sd. Statistical significance was determined by unpaired t-test, ****p < 0.0001. c) DNA content quantification in fresh tendon versus TG (n= 5). d) Pie charts summarizing the relative abundance of major ECM proteins/collagens identified in PC and TG. e) Pathway enrichment analysis of proteins differentially represented between TG and PC. Differential abundance was calculated from log2-transformed protein intensities using Welch’s t-test followed by Benjamini–Hochberg correction. Proteins with adjusted p < 0.05 were used for enrichment analysis. Pathways are plotted by fold enrichment and −log10 (adjusted p value), with bubble size indicating gene count.

### 2.2. TG shows injectability, thermoresponsive gelation, tissue retention, and enzymatic biodegradation

Upon injection into target tissue, the TG must undergo rapid gelation to ensure the retention and confinement of embedded cells. To evaluate this, the temperature-dependent gelation of TG was first confirmed by visual observation of flow behavior before and after incubation at 37°C. At room temperature, the TG remained in a liquid state, demonstrating clear flow upon tilting the vial. In contrast, after incubation at 37°C, the TG rapidly transitioned into a stable gel (**Fig. 2a**). The thermo-responsive behavior of the TG was further quantified via rheological temperature ramps from 20 to 40°C. The storage moduli (G’) for all concentrations increased with temperature and reached a plateau between 35–37°C. Notably, higher concentrations (15 and 20 mg/mL) reached significantly higher plateau moduli, 325±39.7 Pa and 4134±67.2 Pa respectively, compared to lower concentrations (5 and 10 mg/mL), which reached only 4±2.2 Pa and 35±20.1 Pa (**Fig. 2b**). To further characterize material stability, strain-and frequency-dependent rheological profiles were recorded. Tan δ values (G’’/G’), representing the ratio of viscous to elastic rheological properties, remained low and constant for the 15 and 20 mg/mL groups across physiological shear strains (up to 20%) and frequencies (up to 200 rad s^-1^). Conversely, the 5 and 10 mg/mL groups exhibited a rapid increase in tan δ with increasing strain and frequency, suggesting material instability (**Fig. 2c, d**). Consequently, the 15 and 20 mg/mL concentrations were selected for subsequent characterizations.

**Figure 2.**
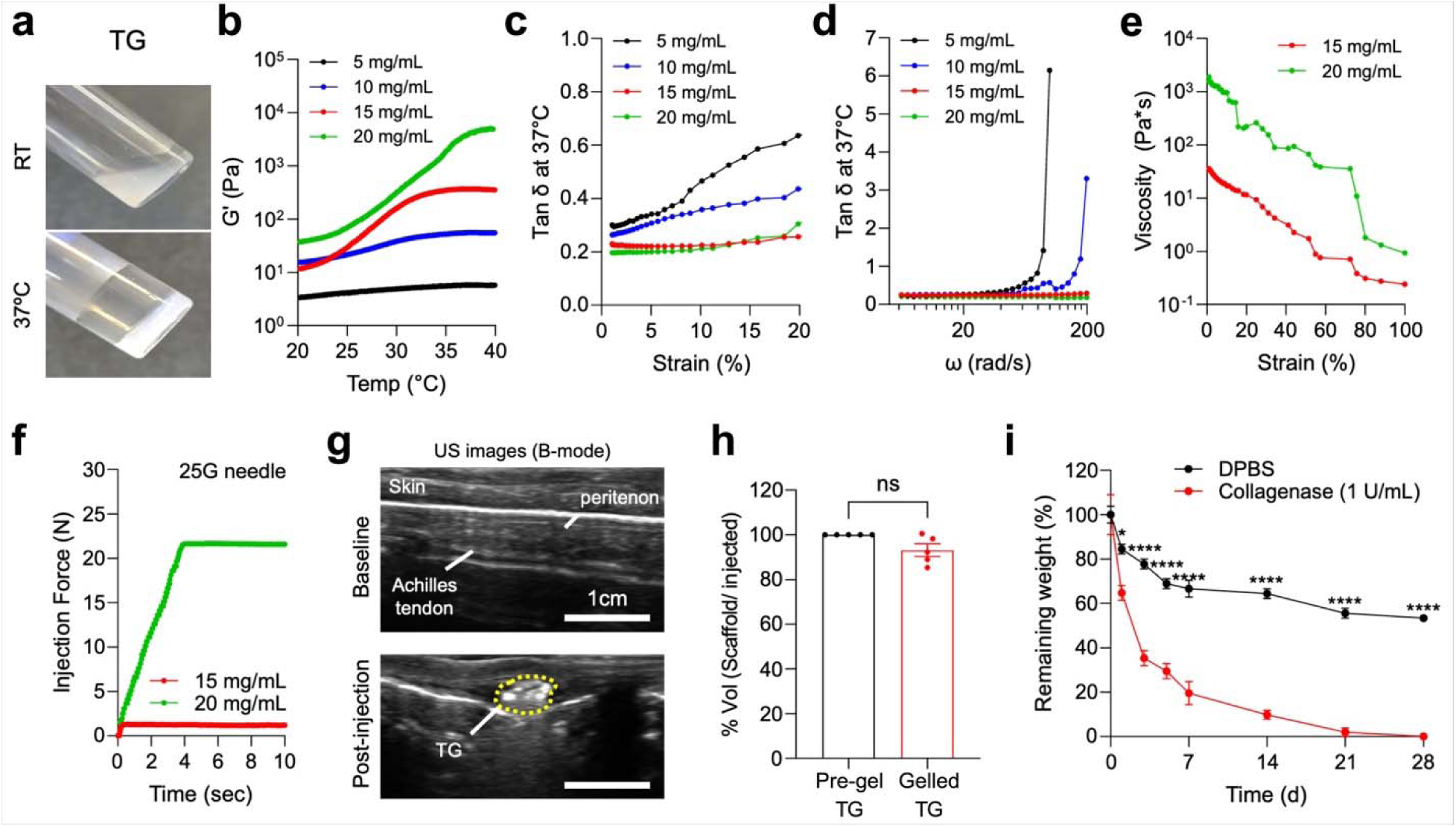
Biomechanical characterization of TG. **a)** Thermoresponsive gelation of the TG. The pre-gel solution is at a liquid state at room temperature but solidifies after incubation at 37°C. **b)** Representative temperature-dependent rheological properties of TG at various concentrations (5–20 mg/mL). Storage moduli (G’) were measured with increasing temperature at a constant frequency of 10 rad s^-1^. **c)** Representative measurements of the loss tangent (Tan δ) at 37°C for different TG concentrations (5–20 mg/mL) at a constant frequency of 10 rad s^-1^. Shear strain ranges from 0 to 20%. **d)** Representative measurements of tan δ at 37°C for various TG concentrations (5–20 mg/mL) across a physiological frequency range (0–200 rad s ¹). **e)** Representative viscosity flow curves of 15 and 20 mg/mL TG with increasing strain (0–100%). **f)** Time-dependent injection force curves of 15 and 20 mg/mL TG injected through a 25-gauge needle, illustrating the compression force buildup associated with the 20 mg/mL concentration. **g)** Representative sagittal ultrasound images before (top) and after (bottom) TG injection into ex vivo porcine Achilles tendon. Scale bar, 1 cm. **h)** Comparison between the injected volume of pre-gel and the calculated volume of the gelled TG after incubation at 37°C (n= 5). **i)** In vitro degradation of TG in DPBS and collagenase-supplemented buffer, demonstrating accelerated enzymatic degradation of TG (n= 3). Data are presented as mean ± sd. Statistical significance between two groups was assessed using an unpaired two-tailed t-test. Significance levels are indicated as follows: ns, not significant; *p < 0.05 and ****p < 0.0001.

Next, we evaluated the injectability and clinical ease-of-use of the TG formulations. Both 15 and 20 mg/mL TG concentrations exhibited characteristic shear-thinning behavior of ECM-derived gels, with viscosity decreasing significantly as the shear rate increased (**Fig. 2e**). This non-Newtonian property is critical for ensuring hand-injectability and providing an adequate operational window during clinical procedures (**Fig. 2e**) [26–29]. To quantify this, injection force (extrusion force) measurements were performed using a 25-gauge needle. The 15 mg/mL TG was easily injectable, requiring a force of only 1–2 N. In contrast, the 20 mg/mL TG reached a force of approximately 22 N within 4 sec, which exceeds the ideal range for manual hand injection (5–20 N). Consequently, the 15 mg/mL concentration was identified as the most mechanically suitable formulation for our application [30] (**Fig. 2f**).

To demonstrate the feasibility of minimally invasive delivery, a percutaneous needle injection into *ex vivo* porcine Achilles tendons was performed under ultrasound (US) guidance.

US imaging was conducted before and after TG injection, followed by incubation at 37°C. Post-injection images revealed effective localization and *in situ* retention of the TG within the peritenon (**Fig. 2g**). Successful retention at the injection site was further validated by comparing the initial volume of the injected pre-gel solution with the final volume of the gel as calculated from the US images. No significant difference was observed between these volumes, suggesting minimal loss of the injected TG (**Fig. 2h**).

Considering its exogenous nature, the biodegradability and biocompatibility of the TG is a critical factor for clinical translation. Elevated levels of collagenase at inflammatory tendinopathy sites are expected to accelerate the degradation of the TG, given its collagen-based composition [31]. To evaluate this, the degradation profile of the TG was monitored at 37°C in both the presence and absence of collagenase (1 U/mL Collagenase I). Under control condition (DPBS), TG exhibited a gradual degradation profile, losing 47% of its initial weight over the 28-day observation period. In contrast, the addition of collagenase significantly accelerated the process; the TG lost 35% of its initial weight within the first 24 hours and was almost completely degraded by Day 21 (**Fig. 2i**). This rapid mass loss is primarily attributed to the enzymatic cleavage of the collagen fibers within the TG matrix [32]. To evaluate the biocompatibility of the hydrogel, an in vitro cytotoxicity assay was performed using the TG eluates across a gradient of dilution factors (1 to 128-fold) in accordance with ISO 10993 guidelines. Quantitative viability analysis revealed that cell survival remained stable near 100% relative to the control group across all dilution factors, indicating the absence of leachable toxic components within the scaffold matrix (**Fig. S3**).

### 2.3. TG supports iTenocyte distribution, interface migration, and sustained collagen release *in vitro*

The spatial distribution of iTenocytes within TG was assessed over time *in vitro*, with quantification of collagen release following iTenocyte incubation in TG versus PC. The iMSCs were transduced with a Scleraxis (SCX) lentiviral GFP reporter to generate SCX-GFP^+^ iTenocytes, as confirmed by representative overlaid phase-contrast and fluorescence images (**Fig. S4a**). In the TG culture assay, iTenocytes embedded within TG demonstrated a distinct interface between the culture plate and the TG region **(Fig. S4b)**. Time-course fluorescence imaging revealed progressive migration of iTenocytes toward and beyond the TG boundary, with increasing SCX-GFP signal detected in regions external to TG from Day 1 through Day 3 and 7 **(Fig. S4b)**. At Day 7, samples were imaged in separate channels for nuclei (blue) and SCX-GFP (green), and F-actin (red). Merged images demonstrated the presence of iTenocytes and organized actin structures at the TG–plate interface and in adjacent regions **(Fig. S4b)**.

To further assess three-dimensional cell distribution within TG, confocal laser scanning microscopy (CLSM) was performed on Day 3 and Day 7 using immunofluorescent staining for nuclei, F-actin, and SCX (**Fig. S4c**). Volumetric reconstructions showed widespread distribution of nuclei and cytoskeletal structures throughout the TG matrix at both time points. Higher-magnification views confirmed SCX-GFP expression co-localized with F-actin–positive cellular structures at Day 3 and Day 7 (**Fig. S4c**). Notably, cells exhibited increased cell-cell interconnectivity over time. In parallel with these morphological observations, biochemical analysis of collagen release demonstrated time-dependent increases in collagen type 1 (Col1) and collagen type 3 (Col3) from Day 1 to Day 7 under both TG (iTeno-TG) and PC (iTeno-PC) conditions. (**Fig. S4d**). Col1 release was consistently higher in TG compared with PC at all time points, whereas Col3 increased over time in both conditions with comparable magnitudes.

### 2.4. TG promotes tenogenic maturation and ECM-associated transcriptional remodeling in iTenocytes

Bulk RNA sequencing was performed to quantify gene-level changes in iTeno-TG and iTeno-PC over time and to identify pathways enriched in association with these transcriptional differences (**Fig. 3a-c**). iTenocytes cultured in TG exhibited increased expression of key tenogenic regulators at Day 7 compared with Day 3, including SCX and MKX (**Fig. 3a**). Elevated expression at Day 7 was also observed for tendon matrix–associated genes, including DCN, BGN, TNC, THBS4, POSTN, and DDR2 (**Fig. 3a**). KEGG pathway enrichment analysis comparing Day 7 versus Day 3 iTeno-TG samples identified significant enrichment in pathways related to cytoskeleton organization in muscle cells, focal adhesion, and ECM–receptor interaction (**Fig. 3b**). Gene ontology biological process (GO BP) analysis further identified significant enrichment of terms related to extracellular matrix organization and collagen fibril organization (**Fig. 3c**).

**Figure 3.**
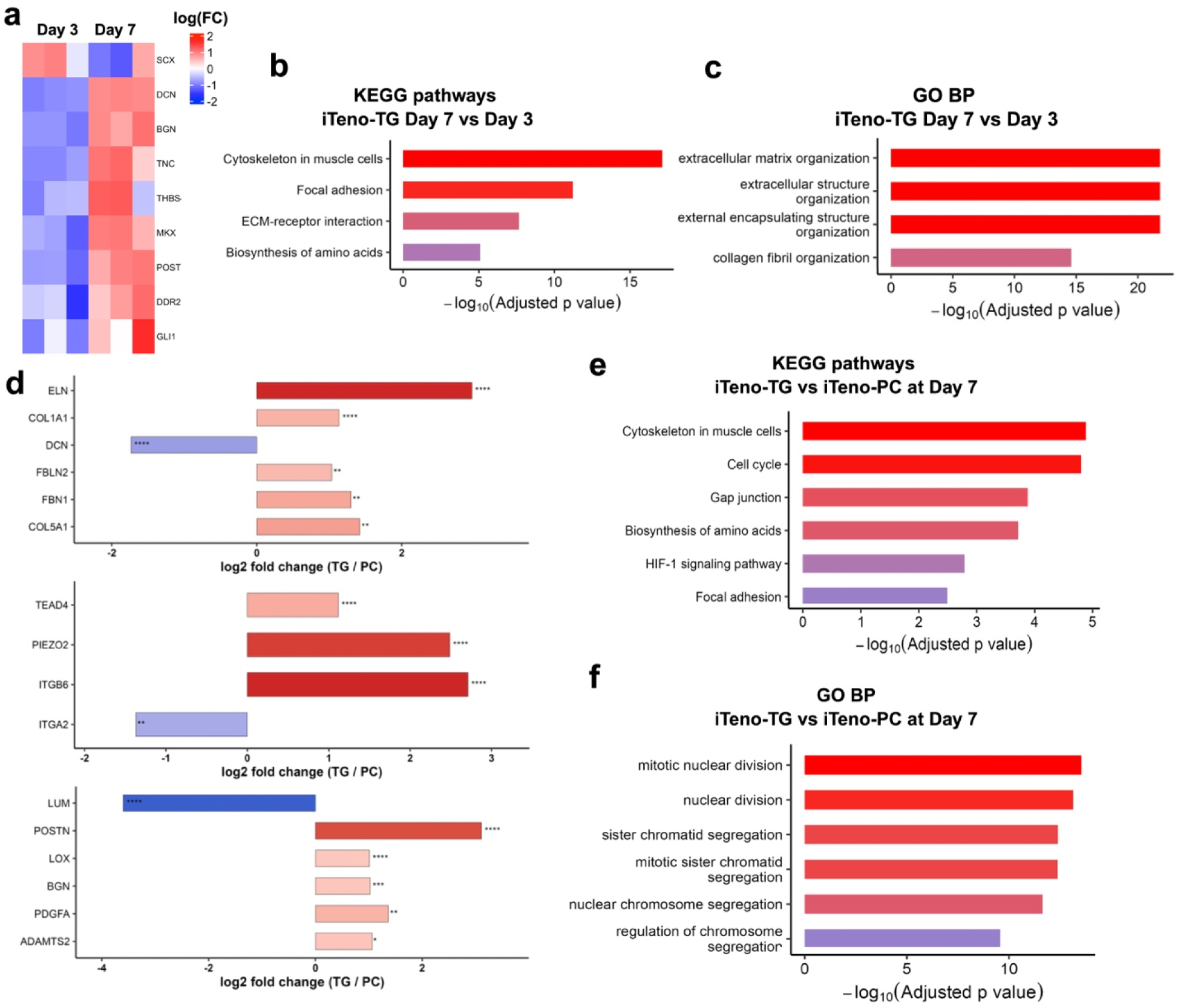
Bulk RNA sequencing reveals time-dependent transcriptional changes in iTenocytes cultured in TG (iTeno-TG) and matrix-dependent transcriptomic differences between TG and PC. **a)** Heatmap of differentially expressed genes in iTenocytes cultured in TG for 3 and 7 days, respectively. **b)** KEGG pathway enrichment analysis for iTeno-TG at Day 7 versus Day 3. **c)** Gene Ontology Biological Process (GO BP) enrichment analysis for iTeno-TG at Day 7 versus Day 3. **d)** Selected gene-level comparisons categorized into tenogenic genes, progenitor cell marker genes, ECM-related genes, and mechanoresponsive genes. **e)** KEGG pathway enrichment analysis of transcriptomic differences between iTeno-TG and iTeno-PC at Day 7. **f)** GO Biological Process enrichment analysis of transcriptomic differences between iTeno-TG and iTeno-PC at Day 7. Statistical significance was determined using differential expression testing with Benjamini–Hochberg correction for multiple comparisons. KEGG and GO BP enrichment analyses were performed using differentially expressed genes, with pathway significance shown as −log_10_ (p-adj). For gene-level comparisons, statistical significance is indicated as: *p < 0.05, **p < 0.01, ***p < 0.001, and **\*\***p < 0.0001.

Genomic differences between iTeno-TG and iTeno-PC following 1 week of maturation were also identified (**Fig. 3d-f**). Gene-level comparisons demonstrated significantly higher expression in iTeno-TG for genes encoding structural ECM components and elastic fiber-associated proteins, including ELN, FBN1, COL1A1, and COL5A1 (**Fig. 3d, Tenogenic genes**). Within ECM-related groups, tendon repair-associated genes such as POSTN, lysyl oxidase (LOX), the collagen processing enzyme ADAMTS2, and the proteoglycan BGN were also upregulated in iTeno-TG compared with iTeno-PC **(Fig. 3d, ECM-related genes)**. In addition, the mechanosensitive genes PIEZO1 and ITGB8 were upregulated in iTeno-TG, whereas ITGA2 expression was higher in iTeno-PC **(Fig. 3d, Mechanoresponsive genes)**. Pathway enrichment analysis of differential gene expression between iTeno-TG and iTeno-PC identified significant KEGG pathways, including cytoskeleton organization in muscle cells, cell cycle, gap junction, biosynthesis of amino acids, focal adhesion, and ECM–receptor interaction **(Fig. 3e)**. GO BP enrichment analysis was dominated by cell division–associated processes, including mitotic nuclear division, nuclear division, chromosome segregation, and nucleosome assembly **(Fig. 3f)**.

### 2.5. iTeno-TG exhibits a tendon-enriched and ECM-remodeling proteomic signature

To correlate RNA expression with functional protein expression, quantitative proteomics was performed to compare relative protein abundances between iTeno-TG and iTeno-PC. Unsupervised hierarchical clustering of differentially abundant proteins clearly segregated the iTeno-TG and iTeno-PC samples, revealing distinct and contrasting protein abundance profiles between the two groups **(Fig. 4a)**. Tendon-related proteins were more abundant in iTeno-TG relative to iTeno-PC **(Fig. 4b)**, including THBS4, DCN, COL5A1, and COL6A2 (Tendon-related proteins). Within the ECM remodeling category, iTeno-TG exhibited higher levels of TIMP3 and SERPINE2, with increased abundance of CTSD and ADAMTSL1 (ECM remodeling proteins). In the proteoglycan category, iTeno-TG showed elevated levels of LUM, ASPN, TGFBI, DCN, BGN, and GPC1 (Proteoglycans). For mechanoresponsive proteins, iTeno-TG showed higher abundance of ITGA2 and LIMK1 whereas FLNB and FBLIM1 were less abundant in iTeno-TG relative to iTeno-PC (Mechanoresponsive proteins).

**Figure 4.**
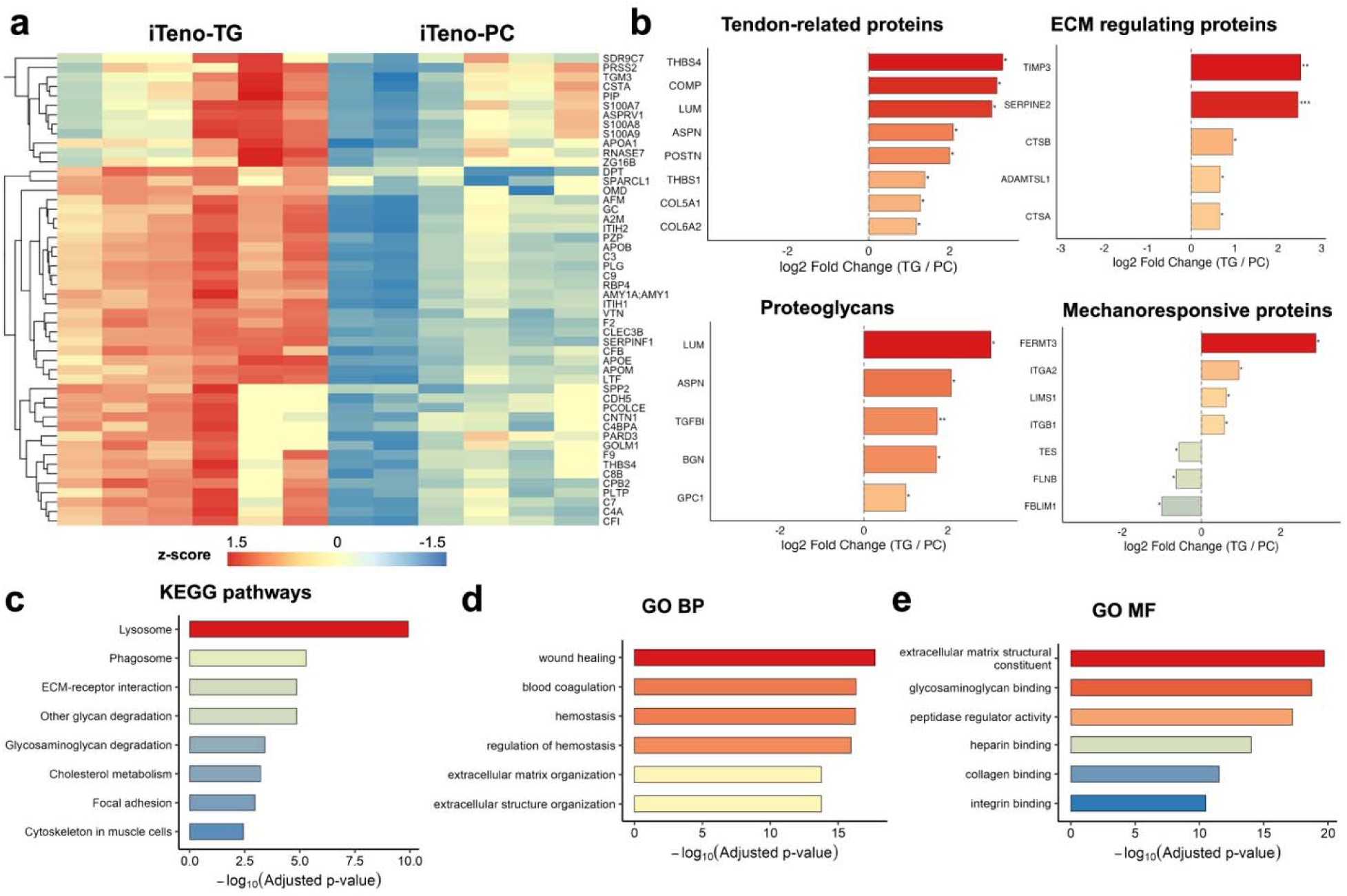
Proteomic comparison of iTeno-TG and iTeno-PC. **a)** Heatmap of differentially abundant proteins in iTenocytes across TG and PC conditions. **b)** Selected protein-level differences presented as log fold change (TG over PC), grouped into tendon-related proteins, ECM remodeling proteins, proteoglycans, and mechanoresponsive proteins. **c)** KEGG pathway enrichment analysis based on proteins differentially abundant between TG and PC. **d)** GO BP and **e)** GO MF enrichment analysis for TG versus PC differentially abundant proteins. Differential protein abundance between TG and PC was assessed using adjusted p-values, with significance levels indicated as follows: *p-adj < 0.05, **p-adj < 0.01, ***p-adj < 0.001, and ****padj < 0.0001. KEGG and GO enrichment analyses were performed using significantly TG-enriched proteins, with enrichment significance shown as −log_10_(Benjamini– Hochberg adjusted p-value).

Pathway enrichment analysis based on differentially abundant proteins identified KEGG pathways including ECM–receptor interaction, focal adhesion and cytoskeleton in muscle cells **(Fig. 4c).** GO BP enrichment highlighted terms such as wound healing, and hemostasis (**Fig. 4d**). Gene ontology molecular function (GO MF) analysis revealed enrichment in extracellular matrix structural constituent, glycosaminoglycan binding, heparin binding, collagen binding, integrin binding, and peptidase inhibitor/regulator activity **(Fig. 4e)**.

### 2.6. iTeno-TG demonstrates localized delivery and short-term cell retention in a rat Achilles tendon defect

To evaluate short-term in vivo retention of implanted cells, iTeno-PC and iTeno-TG were delivered into a partial window defect in the rat Achilles tendon. Longitudinal tracking was performed using an in vivo imaging system (IVIS) on days 3 and 10, followed by histological and immunofluorescence analyses (**Fig. 5a**). IVIS imaging demonstrated localized signal at the delivery site at both day 3 and 10 in the iTeno-PC and iTeno-TG groups, indicating persistence of the constructs during the observed post-implantation period (**Fig. 5b** and **Fig. S5**). The total radiant flux of DiD-labeled iTenocytes was quantified within a consistent region of interest drawn over the tendon defect zone (**Fig. 5c**). To provide a longitudal measure of DiD-labeled iTenocyte persistence within the defect region, the day 10 signal was normalized to the corresponding day 3 signal for each sample and presented as fold change (**Fig. 5d**). These analyses revealed that TG exhibited a cell retention capability comparable to that of the commercially available collagen hydrogel (PC). Histological evaluation using H&E staining further confirmed the presence of cell-containing material within the defect zone at both time points. In the iTeno-PC group, the implanted material remained clearly identifiable within the defect and maintained direct contact with the surrounding native tendon tissue (**Fig. 5e, f**). A similar pattern was observed in the iTeno-TG group, where the implanted gel and associated cells were evident within the defect site and adjacent to the host tendon at both time points (**Fig. 5g, h**). Collectively, these findings demonstrate effective early retention of both iTeno-PC and iTeno-TG within the partial Achilles tendon defect over the 10-day observation period.

**Figure 5.**
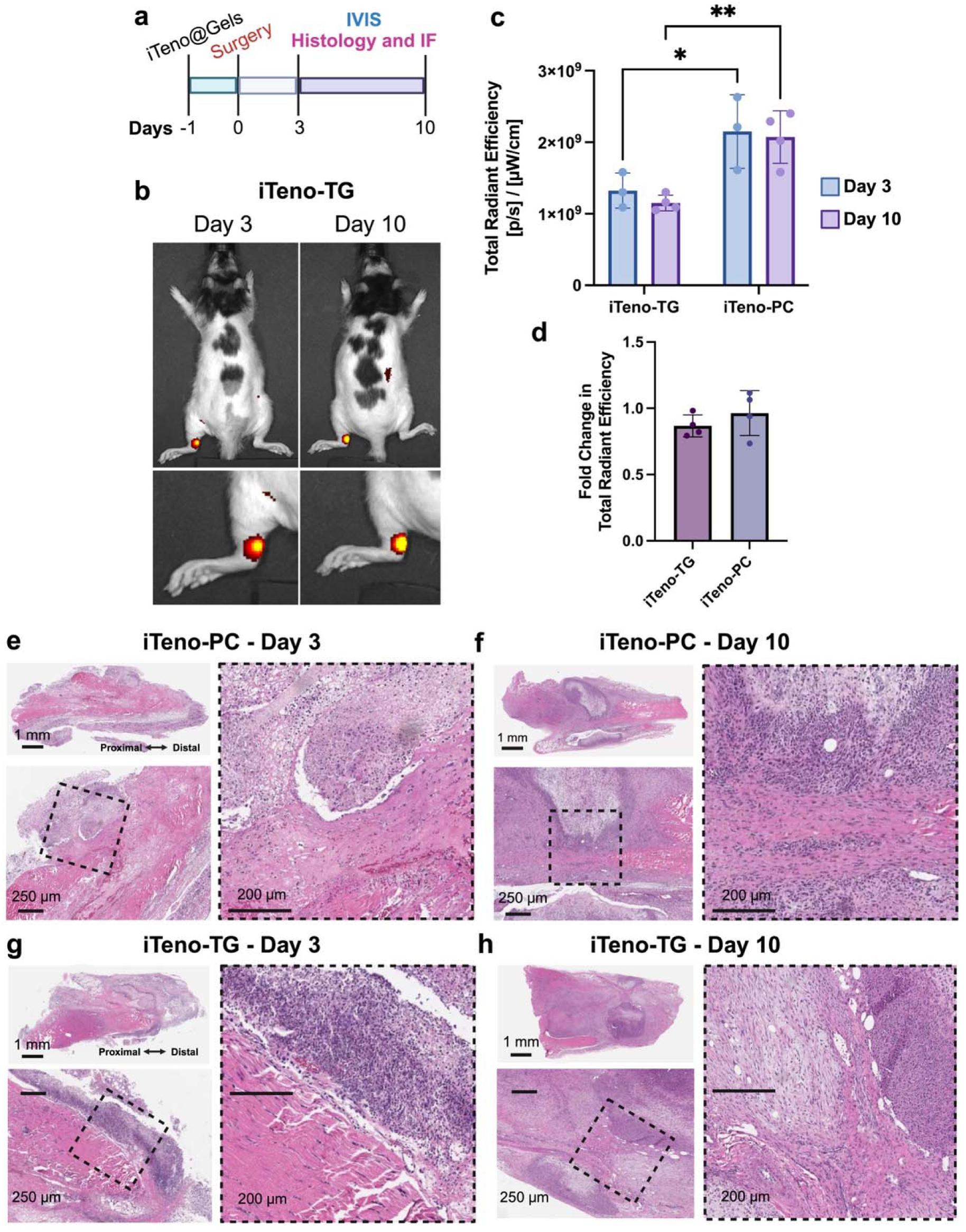
In vivo retention study of iTeno-PC and iTeno-TG in a rat Achilles tendon partial defect model. **a)** Experimental timeline of the pilot study. iTeno-PC or iTeno-TG was implanted into a partial rat Achilles tendon defect. IVIS was performed on days 3 and 10, followed by histology and immunofluorescence. **b)** Representative IVIS images of iTeno-TG at days 3 and 10. **c)** Quantification of IVIS total radiant efficiency within the tendon defect gap at day 3 and 10. **d)** The fold change of total radiant efficiency relative to day 3. Data are presented as mean ± s.d. **e, f)** Representative H&E images of iTeno-PC at days 3 and 10, respectively. **g, h)** Representative H&E images of iTeno-TG at days 3 and 10, respectively. Scale bars: 1 mm for whole area images and 200 μm for close-up images.

Confocal microscopy was used to visualize the implanted cells in relation to the boundaries of the TG construct. High-magnification imaging enabled clear distinction between DiD-labeled iTenocytes and the unlabeled native tendon when positioned in close proximity (**Fig. 6**). At Day 3 post-injury, iTeno-TG was detected on both sides of the native tendon within the full-width biopsy punch defect, which spanned approximately 85% of the tendon width (1.5 mm punch) (**Fig. 6a**). Higher magnification images confirmed the presence of DiD-positive iTenocytes distributed throughout the defect zone. The 50 μm sections demonstrated the presence of both iTeno-TG and host tendon within the same field. By Day 10, iTeno-TG remained localized within the defect site and exhibited a more compact morphology (**Fig. 6b**). Volumetric imaging at this time point showed persistence of the DiD signal within the defect region; however, overall signal intensity was reduced compared to Day 3, suggesting a decrease in the number of detectable labeled cells by Day 10 **(Fig. 6c, d)**.

**Figure 6.**
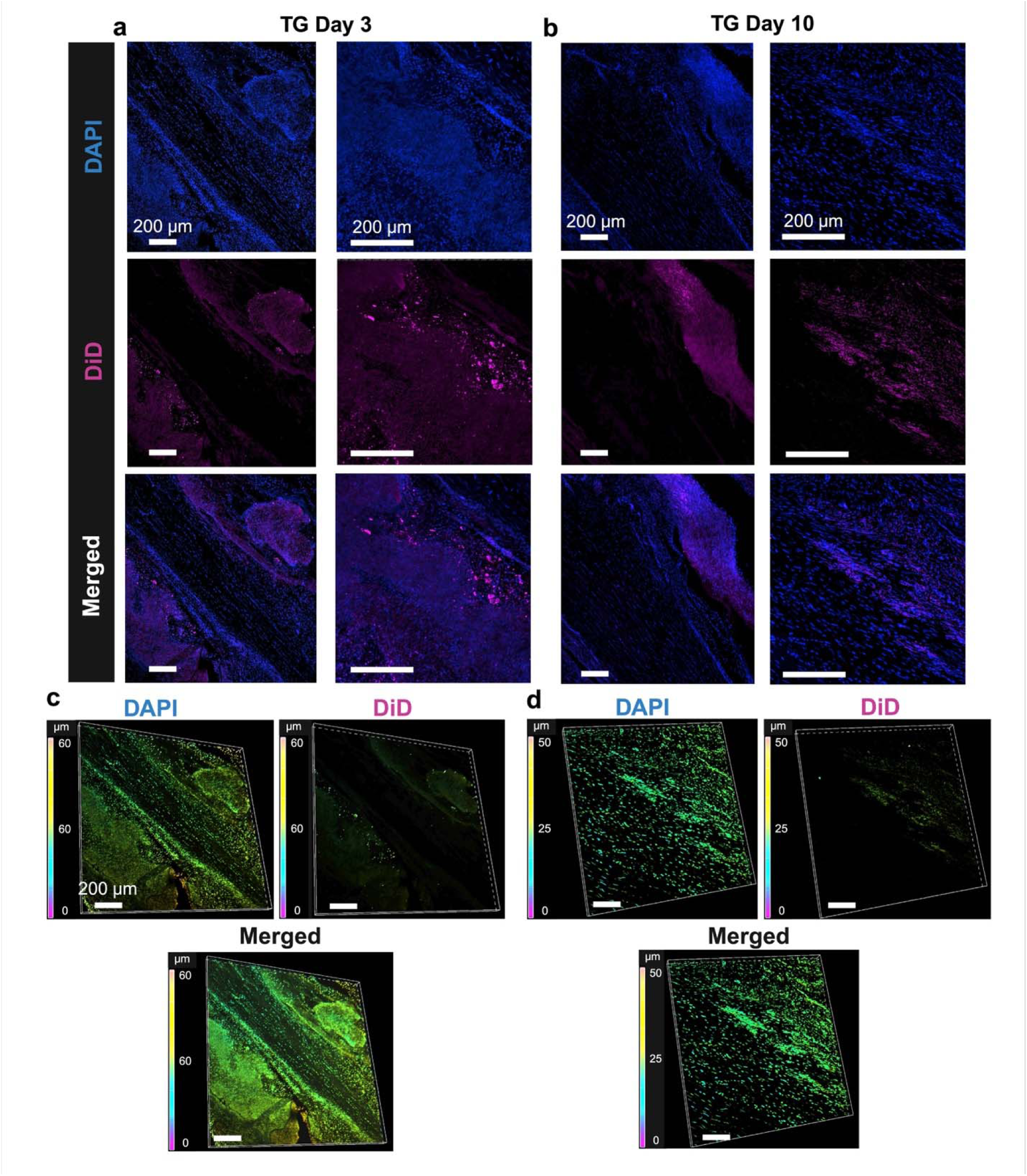
Short term cell retention study of iTeno-TG in rat model. **a)** iTeno-TG was detected along both sides of the native tendon at day 3. **b)** At day 10, iTenocytes remained viable and localized within the defect region. Volumetric images demonstrate the presence of iTenocytes within TG at the defect zone **c)** 3 days post-injury and **d)** 10 days post-injury. Scale bars: 200 μm.

## Discussion

Tendon tissue is characterized by a low metabolic rate and sparse resident cell populations, which severely limits its innate capacity for functional repair. Cell-based therapies have emerged as a promising strategy to supplement the defect site with a viable population of tenogenic cells capable of driving de novo matrix synthesis and orchestrating regenerative signaling. However, the efficacy of this approach is fundamentally constrained by the performance of the carrier at the defect site [33, 34]. As an injectable carrier, TG allows for minimally invasive delivery while conforming to irregular tendon defects. It plays a major role in augmenting tissue architecture rather than limiting spatial regeneration within the tendon [35, 36]. Moreover, carriers must retain cells within the dense, hypovascular tendon environment and provide biochemical cues that preserve the cell phenotype during the early delivery period. While synthetic hydrogels can localize cargo, they often exhibit limited conformity and biological coupling, which can reduce effective cell–host interaction.[37, 38] Consequently, recent studies have utilized composite hydrogel systems to expose transplanted cells to a niche that supports tenogenic maintenance [39–42]. Despite these advances, a robust, clinically practical cell–hydrogel delivery system remains elusive, leaving a major gap in tendinopathy therapies. To address these constraints, we developed an injectable, tendon-derived TG. Designed for minimally invasive placement, TG acts as a biomimetic carrier for iTenocytes. We aimed to improve local cell retention while providing instructive ECM cues to encourage iTenocyte repopulation and maturation within the mechanically active defect environment, where poor cell–tissue contact and cell loss are common barriers to regeneration.

The TG demonstrated a compositionally distinct matrix profile compared with PC, supporting a matrix-mediated mechanism rather than acting solely as an inert cell carrier. Proteomic profiling showed that, while both materials were collagen-rich, the TG retained a broader tendon-associated ECM composition and maintained a collagen profile characterized by greater COL1 representation relative to COL3. Because tendon repair is accompanied by dynamic regulation of collagen isoforms and matrix remodeling [43, 44], this composition supports the use of the TG as a tendon-benchmarked matrix rather than a generic collagen scaffold. Mechanistically, the TG enrichment in proteins associated with ECM–receptor interaction and focal adhesion pathways suggests that the material may provide tendon-relevant adhesive and remodeling cues capable of supporting iTenocyte residence, matrix engagement, and redistribution within the local environment [45, 46]. Consistent with this interpretation, time-course assays showed that iTenocytes localized at the TG–plate interface and progressively migrated beyond the TG boundary from Day 1 to Day 7, indicating that the TG supports cell residence without permanently confining cells. Complementary confocal imaging confirmed three-dimensional cellular distribution within the hydrogel volume. In parallel, the TG showed greater COL1 release over time compared with PC, consistent with increased matrix availability and/or remodeling potential within the TG environment.

Transcriptomic analysis showed that iTeno-TG underwent a coordinated shift from Day 3 to Day 7, characterized by the increased expression of tenogenic regulators (SCX, MKX) and matrix-associated genes (DCN, BGN, TNC). Compared to PC, the TG upregulated genes linked to ECM architecture, remodeling, and mechanosensitive signaling. Interestingly, canonical markers like TNMD were not uniformly different between the TG and PC, suggesting that TG’s advantages lie in controlling how cells adhere, transduce cues, and remodel the 3D environment rather than simply increasing identity markers [47–49]. Consistent with this interpretation, proteomic profiling revealed clear separation between TG and PC conditions. The iTeno-TG demonstrated that the multiple tendon ECM–associated proteins are increased, spanning glycoproteins (e.g., THBS4, COMP), proteoglycans (e.g., LUM, DCN, BGN, ASPN), and collagen-associated components (e.g., COL5A1, COL6A2). The TG also exhibits higher levels of remodeling and protease regulation-related factors, including TIMP3 [50] and SERPINE2 [51] as well as lysosomal proteases CTSD-CTSA [52]. Along with these protein-level differences, functional enrichment highlights pathways linked to ECM–receptor interaction, focal adhesion, and cytoskeleton together with categories related to lysosomal processes. GO enrichment further identifies terms associated with wound healing, coagulation/hemostasis, and ECM-related molecular functions such as extracellular matrix structural constituent and glycosaminoglycan binding. Previously studied in various tissues, a dynamic crosstalk between internal lysosomal activity with external ECM-originating remodeling cues are key factors in regenerative remodeling of the microenvironment [53–55]. Collectively, this suggests that TG culture promotes a profile of matrix assembly and turnover machinery necessary for regenerative remodeling.

Early survival at the defect site is critical for cell-based therapies, as cells must first remain viable before they can populate and contribute to tissue repair [56]. In this context, cytocompatible tissue-derived ECMs have previously been shown to reduce the risk of cell death while providing a more permissive environment than many synthetic or composite materials, which may limit cell permeability and survival [57]. Conversely, the TG offers a tendon-mimetic ECM microenvironment that may simultaneously support early iTenocyte survival, local retention, and iTenocyte maturation after implantation. Among injectable and gel-like systems for tendon repair, most reported platforms primarily function as delivery matrices for cells and/or exogenous bioactive molecules. Although several of these systems support tendon-related repair, their bioactivity often depends on added factors such as PRP [58], TGF-β3 [59], kartogenin [60], extracellular vesicles [61] or mechanical pre-loading of the cell/gel combination [62]. In contrast, TG provides a tissue-derived tendon ECM–based carrier that supports tenogenic activity without requiring additional pro-tenogenic supplementation. To determine whether this effect reflected the TG microenvironment rather than only the intrinsic tenogenic phenotype of SCX iTenocytes, we compared iTeno-TG with iTeno-PC. Although iTenocytes performed well in both carriers, TG further enhanced tendon-related responses relative to the collagen control, suggesting functional synergy between the tendon-derived ECM carrier and the SCX^+^ overexpressing lineage-committed iTenocyte payload. This distinction is important because many reported cell-gel systems use broadly multipotent adipose-derived stem cells [63–65] or immature tendon-derived stem/progenitor cells [66, 67], which may provide regenerative potential but lack the same degree of defined tendon-lineage commitment as iTenocytes. Therefore, iTeno-TG represent a simplified and targeted pro-tenogenic delivery unit, combining a tendon-instructive ECM carrier with a tendon-committed cell source while enabling evaluation of early cell survival, local retention, and implantation efficiency.

iTenocyte delivery from the TG provides evidence of early construct retention rather than definitive proof of long-term iTenocyte survival, migration, or regenerative contribution. DiD labeling and IVIS imaging confirmed persistence of cell-associated signal within the defect, while H&E verified construct localization, but these methods cannot independently resolve cell viability or fate. In addition, the high initial cell loading density may have contributed to gel compaction, contraction, and reduced detectable signal over time. Future studies using genetically labeled iTenocytes will enable more precise tracking of cell survival, localization, and integration.

Taken together, the TG functions as an injectable, conformable, and bio-instructive carrier that may enable minimally invasive delivery and localized retention of therapeutic cells. Within this tendon-derived matrix environment, iTenocytes may further support tenogenic ECM programs and reinforce tendon-lineage commitment. Coupling TG with a lineage-committed iTenocyte payload therefore provides a defined carrier–cell unit to test whether simultaneous improvement of cell delivery, retention, and tendon-relevant molecular signaling can address key implantation barriers in tendon cell therapy.

### Experimental Section

#### Porcine tendon decellularization and TG production

To prepare TG, freshly harvested porcine Achilles tendon from a 6-month-old female Yucatan minipig (Sinclair Bio Resources LLC, Windham, ME, USA) was decellularized according to previously reported protocols with modifications [25, 68]. The tendon was cut into small pieces and washed in 1× PBS under magnetic stirring for 24 h, followed by washing in distilled water (DW) for an additional 24 h to remove tissue residues and contaminants. Decellularization was performed by incubating the tissue in 1% (v/v) Triton X-100 (Sigma-Aldrich, St. Louis, MI, USA) in 100 mM Tris-HCl buffer (pH 8) for 24 h. The samples were then rinsed three times with DW and incubated in 2% sodium deoxycholate (Sigma-Aldrich) overnight. The samples were again rinsed three times with DW and washed overnight to remove residual detergent. The tissue was subsequently incubated in 50 U/mL DNase I (Sigma-Aldrich) in 1 M NaCl solution (Sigma-Aldrich) for 3 h, followed by overnight washing in DW. All buffer changes were performed under sterile conditions in a biosafety cabinet. After processing, the tissues were snap-frozen in liquid nitrogen and lyophilized for 24 h. The lyophilized tissues were solubilized in 1 mg/mL pepsin (Sigma-Aldrich) in 0.1 M hydrochloric acid (Sigma-Aldrich) at ambient temperature under UV light for 3 days. The solubilized TG was aliquoted and stored at 4°C for short-term use or at-20°C for long-term storage. Prior to use, 800 µL of TG was transferred into a microcentrifuge tube on ice, followed by the addition of 120 µL 10× MEM (Thermo Fisher), 200 µL TGM, and 80 µL 1 N NaOH (Sigma) to neutralize. The components were mixed with chilled wide-bore 1 mL pipette tips until a uniform solution was achieved.

#### Total proteomics of TG and PC hydrogels

TG and PC (Purified collagen from rat tail collagen, Sigma) hydrogel samples (n=6) were collected after gelation in incubator overnight and lysed in S-Trap high-recovery lysis buffer containing 5% SDS, 8 M urea, and 100 mM glycine at pH 8.5. Lysates were incubated on ice for 30 min, sonicated, and centrifuged at 12,000 × g for 10 min at 4°C, and protein concentration was determined by BCA assay following the supplier’s protocol (Thermo Fisher). One hundred micrograms of protein from each sample were adjusted to same volume, reduced with 1 M DTT for 15 min at 37°C, and alkylated with 1 M iodoacetamide for 30 min in the dark. Samples were then acidified with 12% phosphoric acid (∼1.2% final) and mixed with S-Trap binding buffer containing 100 mM TEAB in 90% methanol. Proteins were processed on S-Trap micro columns, washed three times with excess binding buffer, and digested on-column with trypsin in 50 mM TEAB for 1 h at 47°C. Peptides were sequentially eluted with 50 mM TEAB, 0.1% formic acid in water, and 0.1% formic acid in 50% acetonitrile, then collected for mass spectrometry-based total proteomic analysis. The Data Independent Acquisition (DIA) analysis was performed on an Orbitrap Astral (Thermo Scientific) mass spectrometer interfaced with an EASY-Spray™ nano-electrospray ionization source (Thermo Scientific, ES081) coupled to Vanquish Neo ultra-high-pressure chromatography system with 0.1% formic acid in water as mobile phase A and 0.1% formic acid in acetonitrile as mobile phase B. Peptides were separated at an initial flow rate of 1.2 µL/minute and a linear gradient of 4-9% B for 0-2 minutes, 9-25% B for 2-18 minutes, 25-35% B for 18-27 minutes. The flow rate was then increased to 3 µL/min and the column was then flushed with 35-99% B for 0.4 minutes then held at 99% B for 0.5 minutes, decreased to 5%, and the zebra wash flushing was repeated two more times. The column used was PepSep C18 15cm x 150 µm, 1.5µm (Bruker, P/N: 1893474). Source parameters were set to a voltage of 2300 V and a capillary temperature of 280°C. MS1 scan range was set to 360-1200 m/z and MS1 resolution was set to 240,000 with an AGC target set to ‘Custom’ and a normalized AGC target set to 500%. RF Lens was set to 40% with maximum injection time set to 10ms. Precursor mass range was set to 380-980 m/z with 199 non-overlapping data independent acquisition precursor windows of size 3 m/z. MS2 scan range was set to 150-2000 m/z and normalized HCD collision energy set to 30%. Maximum injection time was set to 2.7ms with AGC target set to ‘Custom’ and normalized AGC target set to 500%. All data is acquired in profile mode using positive polarity. After collecting output, the total proteomic profiles of TG and PC hydrogels were compared using three replicates per group in R (v4.4.2). Protein abundance values were log2-transformed, and mean expression levels were calculated for each group to determine differential protein abundance as log2 fold change (TG − PC). The most altered proteins were visualized by hierarchical clustering heatmaps and fold-change heatmaps. Relative protein composition, including collagen subtype distribution, was further summarized as within-group percentages. For visualization of enriched KEGG pathways, the top-ranked pathways were selected based on fold enrichment. Adjusted p values were transformed as −log10(adjusted p value) for graphical display. Dot plots were generated with fold enrichment on the x-axis, pathway names on the y-axis, point size representing gene count, and color representing significance. Bar plots were generated similarly, with bar fill indicating the transformed adjusted p value.

#### Characterization of rheology, injectability, biodegradability and cytotoxicity of the TG

The rheological properties of TG samples were measured using a rheometer (HR10, TA Instruments, New Castle, DE, USA). Temperature sweeps were performed using a 20 mm cone geometry at a strain of 1% and a frequency of 10 rad/s. The gap was adjusted to fill the measurement volume. Oscillation and frequency sweeps were conducted using an 8 mm sandblasted parallel plate geometry at 37°C, with a strain of 1% and a frequency of 10 rad/s, respectively. Fully crosslinked gels were punched (8 mm diameter), loaded onto the rheometer, and measured at a gap of 1500 μm. The storage modulus (G′), loss tangent (tan δ), and complex viscosity were analyzed. The experiment was conducted with a sample size of n= 3.

Injectability was evaluated using a mechanical tester (Instron 5942, Instron, Norwood, MA, USA). TG gels at concentrations of 15 and 20 mg/mL were loaded into 1 cc Luer-Lok™ syringes (Becton, Dickinson and Company, Franklin Lakes, NJ, USA) and injected through a 25-gauge needle (Becton, Dickinson and Company) at a constant flow rate of 1 mL/min. The injection force was recorded using Bluehill version 3 software (Instron).

Ultrasound (US) images were acquired before and after percutaneous injection of 100 µL of TG pre-gel solution *ex vivo* using a 25-gauge needle and a portable US device (T3200 equipped with 15L4 linear transducer; Terason, Burlington, MA, USA). Following injection, the tissues were incubated at 37°C for 5 min to induce gelation of TG. The volume of the formed TG scaffold was then calculated using the acquired US images. The experiment was conducted with a sample size of n = 5.

For *in vitro* degradation studies, equal volumes of gelled TG samples were prepared by punching cylindrical constructs (1 cm diameter, 2 mm height). These hydrogel samples were placed into individual 5 mL Eppendorf tubes (Eppendorf, Hamburg, Germany) containing 3 mL of 1× PBS with or without 1 U/mL collagenase type I (Sigma-Aldrich), followed by incubation at 37°C under gentle agitation (100 rpm). At predetermined time points (0, 1, 3, 5, 7, 14, 21, and 28 days), TG samples were collected, gently washed three times with DW, and lyophilized. The lyophilized samples were weighed, and the percentage of remaining mass was calculated relative to the initial weight at day 0. The experiment was conducted with a sample size of n = 3.

For cytotoxicity test of TG, 3×10^4^ cells/well of L929 fibroblasts (ATCC, Manassas, VA), primary tenocytes, and mesenchymal stem cells (ATCC) were seeded in 96-well tissue culture plates and incubated overnight in a 37°C, 5% CO_2_ humidified chamber. Following incubation, the culture medium was removed, and the cells were treated with 100 µL of fresh growth medium containing serially twofold diluted TG hydrogel eluates ranging from 1 to 128 dilution factors, prepared in accordance with ISO 10993-5. The cells were incubated with the extracts for 24 h, after which each well was replenished with 100 µL of fresh growth medium and 10 µL of WST-1 reagent, followed by a 2 h incubation period. The absorbance of the samples was subsequently measured using a microplate reader at 450 nm and the percentage of viable cells in each treated well was calculated relative to the untreated negative control group.

#### Generation of SCX^+^ iTenocytes from iPSC-derived mesenchymal cells

Human iPSCs were acquired from the Cedars-Sinai iPSC Core Facility and differentiated into induced mesenchymal stem cells (iMSCs) using our previously reported protocol [69]. The resulting iMSCs were then transduced with an SCX-GFP plasmid (OriGene) using a lentiviral vector according to methods previously established by our group [70]. The iTenocytes were generated to overexpress SCX and exhibit persistent nuclear GFP labeling.

#### iTenocyte loading in TG, fluorescence, confocal laser scanning microscopy and collagen release analysis of iTeno-TG

Neutralized TG and PC aliquots were transferred into 200 µL centrifuge tubes and maintained on ice until use. iTenocytes were thoroughly rinsed with PBS and detached using 0.25% trypsin– 0.1% EDTA in Hank’s Buffered Saline Solution for 5 min, followed by one PBS wash. Cells were then resuspended in tendon growth medium (TGM) consisting of DMEM/F12, 1% non-essential amino acids, 1% antibiotic–antimycotic, and 0.004% 1-thioglycerol. The cell suspension (2000 cell/μL gel) was gently mixed with the corresponding gel aliquots on ice using wide-bore pipette tips and transferred to well plates for gelation. Constructs were incubated at 37°C for 10 min to allow complete gelation, after which TGM was added for the first 24 h. On the following day, the medium was replaced with tendon differentiation medium (TDM) composed of low-glucose DMEM, 1% non-essential amino acids, 1% antibiotic–antimycotic, 50 µg/mL L-ascorbic acid, and 0.004% 1-thioglycerol.

To evaluate iTenocyte migration from TG and their 3D spatiotemporal distribution over 7 days, iTeno-TG constructs were prepared in glass-bottom well plates (n = 8). At each time point, samples (n = 2) were rinsed with PBS and fixed in 4% paraformaldehyde prepared in PBS containing 0.02% sodium azide for 15 min at room temperature in the dark. Samples were then permeabilized with Perm/Wash buffer (BD Biosciences) for 30 min at room temperature under 90 rpm orbital shaking. After washing, samples were stained with 0.1 mg/mL phalloidin-iFluor 555 (Abcam) for 30 min at room temperature and counterstained with 0.1 mg/mL DAPI. Live GFP-tagged iTenocytes migrating from TG were imaged without fixation using the EVOS M5000 Imaging System (Thermo Fisher) to monitor cell egress and TG degradation over time. Fixed and stained samples were imaged using a Nikon AX R laser-scanning confocal microscope (Nikon, Japan) to assess 3D cell distribution and cytoskeletal organization.

To quantify collagen I and collagen III released during the incubation period, samples were prepared in plastic-bottom well plates, and half of the culture medium volume was collected from each well at predetermined time points (n = 5). An equal volume of fresh medium was then added back to maintain the original culture volume. Collected aliquots were analyzed in equal volumes using a Collagen I ELISA kit (MyBioSource, MBS1603537) and a Collagen III ELISA kit (MyBioSource, MBS9716169), according to the manufacturer’s instructions.

#### Bulk RNA-seq analysis of iTeno-TG and iTeno-PC

For bulk RNA-seq analysis, iTeno-TG and iTeno-PC samples were cultured in tenogenic differentiation medium (TDM) for 1 week. Following incubation, samples were washed twice with PBS, removed from the culture plates, and transferred to 1 mL microcentrifuge tubes. Samples were lysed in Buffer RLT (Qiagen) and homogenized for 1 min using a pellet pestle. Total RNA was then isolated and purified according to the manufacturer’s instructions using the RNeasy Mini Kit (Qiagen). Purified RNA was stored at −80°C until further use.

RNA-Seq was done at the Cedars-Sinai Medical Center’s AGCT Core. Total RNA was quantified using Qubit RNA High Sensitivity assay (Invitrogen). RNA quality was assessed with Tapestation High Sensitivity RNA Screentape assay (Agilent). RNA was normalized and libraries prepared for sequencing using the SMART-Seq mRNA LP (with UMIs) library preparation kit (Takara Bio). Final libraries were quantified with Qubit 1X dsDNA High Sensitivity assay (Invitrogen). Library fragment size was measured using Tapestation High Senstivity D1000 Screentape assay (Agilent). Libraries were normalized to 2nM and pooled equimolarly for a final sequencing concentration of 130pM. Sequencing was done on the NovaSeq X Plus at 2×100bp with a 1% PhiX library spike-in.

Raw count data were imported from the numreads.tsv matrix and analyzed in R using DESeq2. For the TG3 versus TG7 comparison, the count matrix was subset to include six samples per group (n = 6). A metadata table defining sample condition was generated, and differential expression analysis was performed. DESeq2 results were extracted using the contrast TG3 versus TG7, such that positive log2 fold change (log2FC) values reflected higher expression in TG3 relative to TG7. Genes with adjusted p value (padj) < 0.05 were considered significantly differentially expressed. For sample-level visualization, variance-stabilizing transformation (VST) was applied, and principal component analysis (PCA) was performed using the transformed counts. Selected tenogenic and ECM-related gene sets were extracted and visualized as row-scaled heatmaps. For functional enrichment analysis, genes from the TG3 versus TG7 DESeq2 comparison were divided into upregulated (log2FC > 0.5, padj < 0.05) and downregulated (log2FC <-0.5, padj < 0.05) sets. Gene symbols were converted to Entrez IDs using org.Hs.eg.db, and Gene Ontology (GO) enrichment for Biological Process as well as KEGG pathway analysis, was performed using clusterProfiler. Enrichment significance was assessed using Benjamini–Hochberg correction, with p and q cutoffs of 0.05.

For iTeno-TG vs iTeno-PC, bulk RNA-seq data from TG and PC samples were analyzed in R using DESeq2, with PC set as the reference condition. Differential expression results, including log2 fold change, Wald test p values, and Benjamini–Hochberg adjusted p values, were extracted for TG versus PC comparisons. VST-transformed expression values were used for heatmap generation. Genes were grouped into functional marker panels and filtered using padj < 0.05 and log2FC ≥ 1 for focused visualization by bar plots and row-scaled heatmaps. For enrichment analysis, upregulated and downregulated genes were identified using log2FC thresholds of >0.5 and <−0.5 with padj < 0.05, converted to Entrez IDs, and analyzed by GO and KEGG pathway enrichment using clusterProfiler.

### Total proteomics analysis of iTeno-TG and iTeno-PC

The similar method as mentioned in total proteomics comparison of TG and PC hydrogels was followed. After collecting data, the sample columns corresponding to TG and PC replicates (n=6) were extracted from the proteomics matrix using R (v4.4.2), and protein intensities were log2-transformed prior to analysis. Mean log2 abundance values were calculated for each group, and differential protein abundance was expressed as log2 fold change (log2FC = TG − PC). For visualization, the top 50 proteins ranked by absolute log2FC were displayed as a row-scaled heatmap, and a separate one-column heatmap was generated to show the direction and magnitude of log2FC values. Proteins were also grouped into predefined functional modules, including tendon core proteins, ECM-related proteins, proteoglycans and mechanoresponsive genes for focused comparison of TG and PC protein profiles.

### In vivo cell retention and histological analysis

The iTenocytes were labeled with DiD lipophilic membrane dye (1 µM per 1 M cells, Thermo Fisher) for in vivo tracking. Labeled cells were mixed with TG aliquots on ice at a final density of 10000 cells/µL gel using wide-bore pipette tips. The resulting iTeno-TG constructs were dispensed as 20 µL droplets into 96-well plates, allowed to fully gel for 5 min, and then covered with pre-warmed tendon growth medium (TGM). After 24 h of incubation, samples were transported to the operating room and maintained in a 37°C water bath until implantation.

Male Sprague Dawley rats (>2 months old, Charles River; n = 14) were anesthetized with 4% isoflurane on thermal pads. After shaving the left hind limb, the surgical field was sterilized with 1% hydrogen peroxide in PBS followed by betadine, and a 1 cm skin incision was made. The Achilles tendon was exposed using curved forceps, and a 1.5 mm punch biopsy was used to create a partial-thickness defect involving approximately 75% of the tendon width, without surgical repair. iTeno-TG and iTeno-PC samples were placed into the tendon defect using a spatula to completely fill the injured region. The skin was closed with bioresorbable sutures. Animals were recovered individually with ad libitum access to food and water, and buprenorphine XR (65 mg/kg) was administered postoperatively.

At each of two predetermined post-injury time points, in vivo fluorescence imaging was performed, after which animals were euthanized by CO inhalation followed immediately by lung puncture to confirm death, in accordance with institutional IACUC approval (No. 008586). Achilles tendons were harvested and fixed in 4% paraformaldehyde overnight. Tissues were then paraffin-embedded, and five micron thick sections were obtained. The sections were then re-hydrated and stained with hematoxylin and eosin (H&E) for morphological feature evaluation. Additional sections were analyzed with CLSM. For in vivo fluorescence tracking, animals were anesthetized with isoflurane and imaged using an in vivo imaging system (IVIS) at the designated post-implantation time points (IVIS Lumina XR). Animals were positioned consistently to visualize the operated Achilles tendon region and DiD-associated fluorescence was acquired using the appropriate near-infrared fluorescence settings. Identical imaging parameters were used across samples within each imaging session. Fluorescence signal was quantified using region-of-interest (ROI) analysis. A fixed-size ROI was placed over the tendon defect gap, and average radiant efficiency was recorded for each sample. Contralateral Achilles tendons and/or local background regions were used as fluorescence controls. To evaluate signal maintenance over time, average radiant efficiency values measured within the defect gap were compared between day 3 and day 10 and expressed as fold change, with day 10 signal normalized to the corresponding day 3 signal for each carrier group. Representative IVIS images were displayed using comparable intensity scaling. Contralateral Achilles tendons were used as controls for fluorescence imaging, histological assessment, and immunofluorescence analysis.

## Statistical Analysis

The statistical analysis for rheological studies and collagen release study was performed in Prism v11 (Graphpad) and analyzed with one-way ANOVA. The significant differences were denoted as *p<0.05, **p<0.01, ***p<0.001 and ****p<0.0001. Proteomic intensities were log2-transformed, and differential abundance between TG and PC was calculated as log2 fold change during both hydrogel-only and cell-loaded hydrogel comparisons. Statistical significance for each protein was determined by group-wise comparison followed by Benjamini–Hochberg correction for multiple testing. Proteins with adjusted p < 0.05 and |log2FC| > 0.5 were considered significant. Differences in the proportional distribution of COL1A1, COL1A2, COL3A1, and other ECM components between TG and PC were evaluated using a Pearson chi-square test of independence. Differential expression between TG3 and TG7 was calculated in DESeq2, which generated log2 fold change, Wald test p values, and Benjamini–Hochberg adjusted p values. Genes meeting padj < 0.05 and |log2FC| > 0.5 were used for downstream GO and KEGG enrichment analyses. For TG versus PC proteomics, group differences were tested on log2-transformed protein abundance values using Welch’s t-test. Log2 fold change was calculated as the difference between mean TG and mean PC abundance values, and Benjamini– Hochberg adjusted p values were used to correct for multiple comparisons. Proteins with padj < 0.05 were considered significantly different.

## Acknowledgements

The research was made possible by a grant from the California Institute for Regenerative Medicine (CIRM EDUC4-12751 to AEP; and CIRM DISC0-14350 to DS). The contents of this publication are solely the responsibility of the authors and do not necessarily represent the official views of CIRM or any other agency of the State of California. We thank F. P. S. Guastaldi for generously providing the porcine tissues used to prepare the TG samples.

## Data Availability

Any information required about the data reported in this paper is available from the lead contact upon request.

## Author Contributions

A.E.P., H.K., B.F., and D.S. conceived the idea. A.E.P and H.K. contributed equally to this work. A.E.P., H.K., B.F., and D.S. designed the experiments. H.K. harvested and processed porcine Achilles tendon for TG. A.E.P. and H.K. performed and analyzed the *in vitro* experiments. A.E.P. performed transcriptomic and proteomic analyses. A.E.P. and M.B. performed rat experiments. A.E.P., H.K., G.D., and M.K. wrote the original manuscript. A.E.P., H.K., M.K., B.F., and D.S. revised the manuscript. B.F. and D.S. are principal investigators of the supporting grants.

