## Supplemental Information for "Tendon-derived injectable bio-instructive gel augments tenogenic differentiation of iMSC-SCX cells and extracellular matrix remodeling"

^1^Orthopaedic Stem Cell Research Lab

^2^Board of Governors Regenerative Medicine Institute

^3^Department of Orthopedic Surgery, Beth Israel Deaconess Medical Center, Harvard Medical School, Boston, MA

^4^College of Medicine, Kyung Hee University, Seoul, Republic of Korea

^5^Clinical Research Institute, Kyung Hee University Hospital at Gangdong, Seoul, Republic of Korea

^6^Department of Orthopaedic Surgery, Kyung Hee University College of Medicine, Kyung Hee University Hospital at Gangdong, Seoul, Republic of Korea.

^7^Department of Orthopedics, Cedars-Sinai Medical Center, Los Angeles, CA.

^8^Department of Surgery, Cedars-Sinai Medical Center, Los Angeles, CA.

^9^Department of Biomedical Sciences, Cedars-Sinai Medical Center, Los Angeles, CA.

^10^Biomedical Imaging Research Institute, Cedars-Sinai Medical Center, Los Angeles, CA.

^†^A.E.P. and H.K. contributed equally to this work.

*Co-senior authors

Corresponding Authors’ emails:

Dmitriy Sheyn:

Freedman, Benjamin:

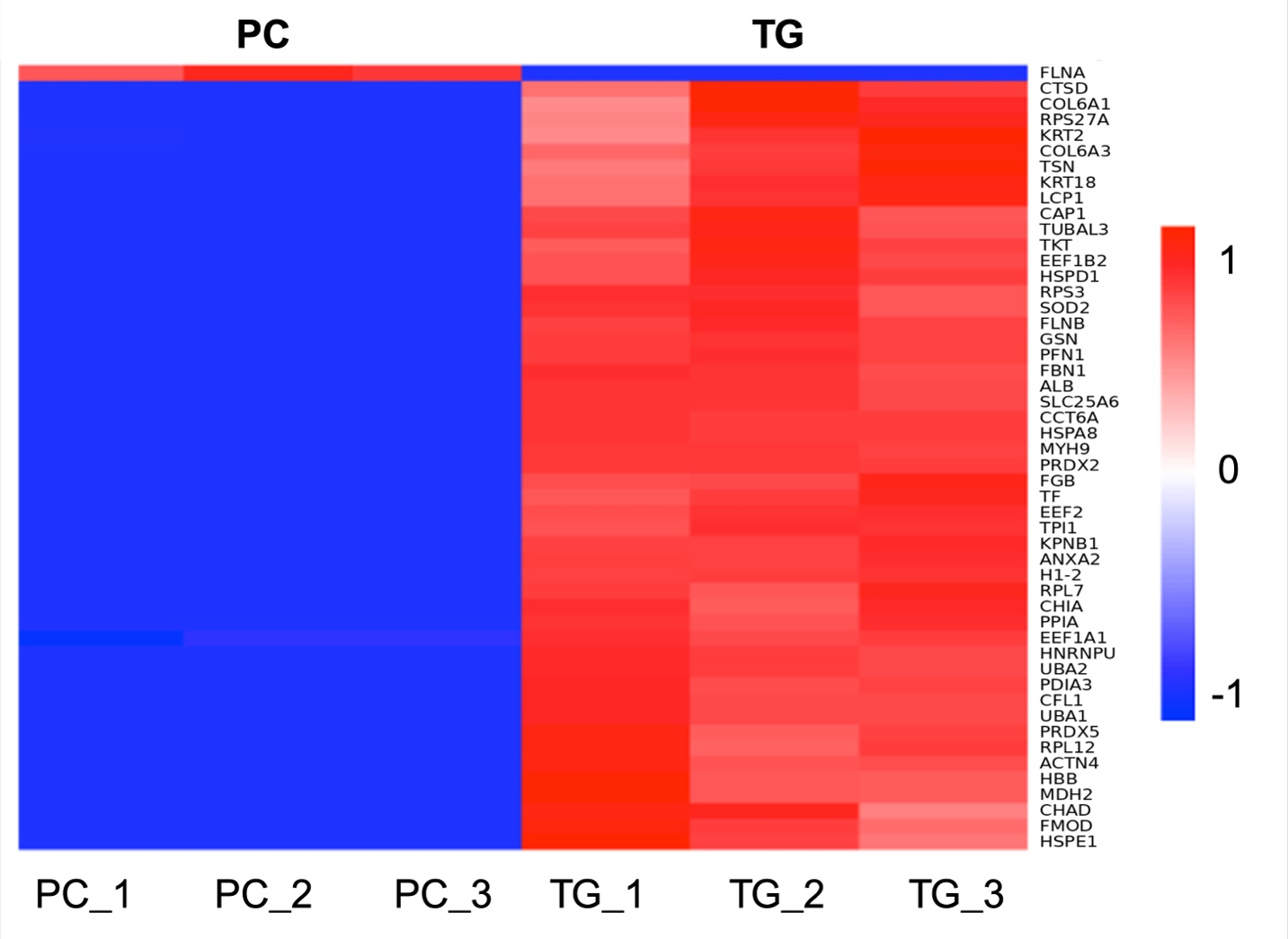

**Figure S1.** Heatmap of proteomic profiles comparing PC and TG samples (PC_1–PC_3; TG_1–TG_3) with hierarchical clustering (relative abundance scale as shown).

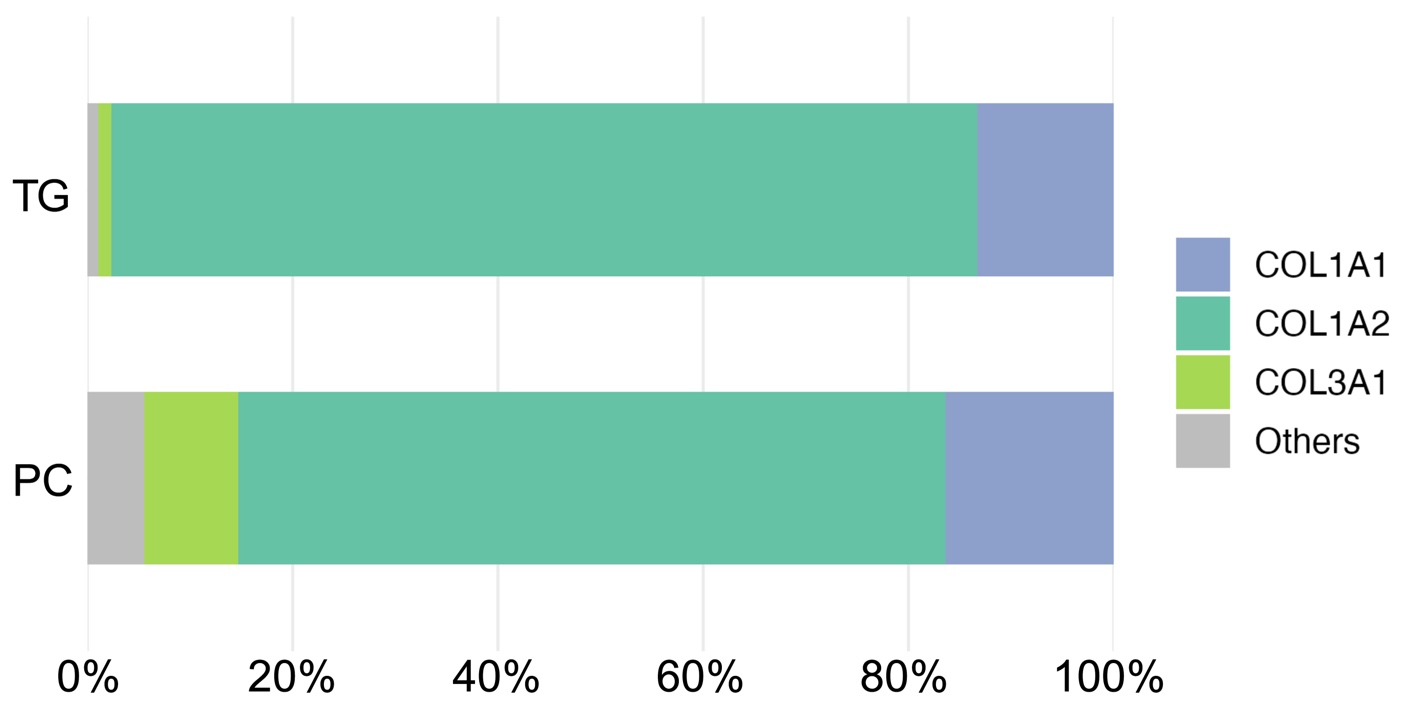

**Figure S2.** Relative abundance of grouped tendon-healing associated collagens was visualized as 100% stacked bars for TG and PC. The relative collagen compositions differed significantly between TG and PC (χ²(3) = 11.94, p = 0.0076).

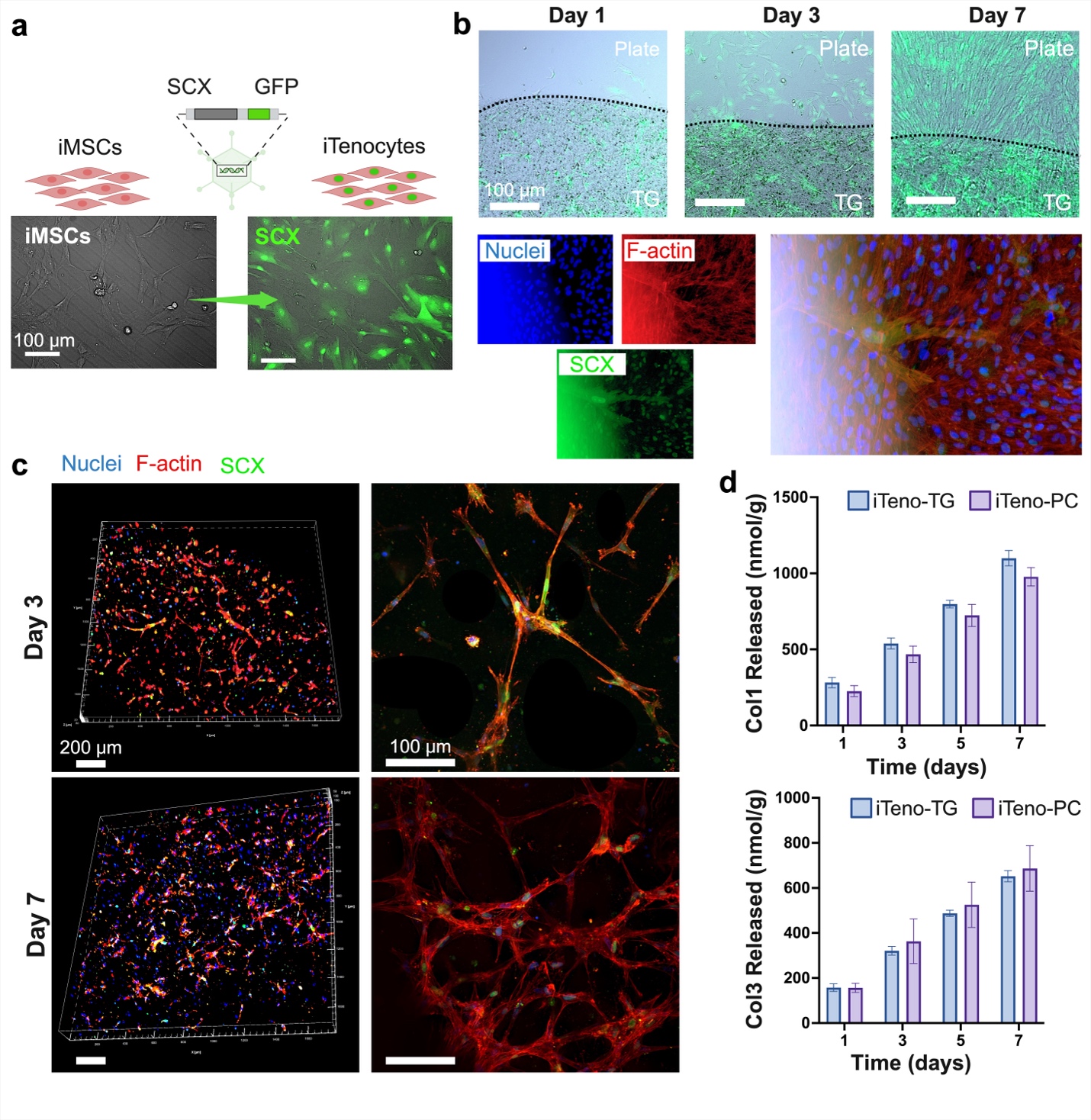

**Figure S3.** **TG functions as an active cell carrier by supporting iTenocytes interconnectivity and migration. a)** Schematic of SCX lentiviral GFP reporter transduction of iMSCs to generate iTenocytes, with representative phase-contrast and fluorescence images. Scale bar, 100μm. **b)** Time-course fluorescence images of iTenocytes cultured in TG, demonstrating dynamic morphological changes at the TG-plate interface over 7 days. Scale bars: 50μm for channel images and 100μm for merged image. **c)** Confocal microscopy images of iTenocytes within TG at Day 3 and Day 7, showing three-dimensional z-stack reconstructions and representative higher-magnification views. Scale bars: 200μm for volumetric views and 100μm for higher-magnification images. **d)** Quantification of accumulative collagen release (nmol/g) for Col1 and Col3 from iTenocyte cultures in TG versus PC over time. Data are presented as mean ± s.d.

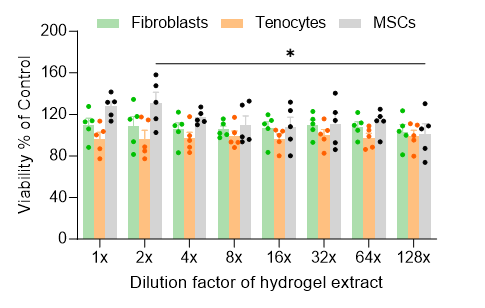

**Figure S4.** Viability of rat fibroblasts, primary rat tenocytes, and human mesenchymal stem cells following 24 h of incubation with TG eluates at various dilution factors (1~128x). Data are mean ± sd. Statistical analysis was performed using two-way ANOVA. * *p* < 0.01.

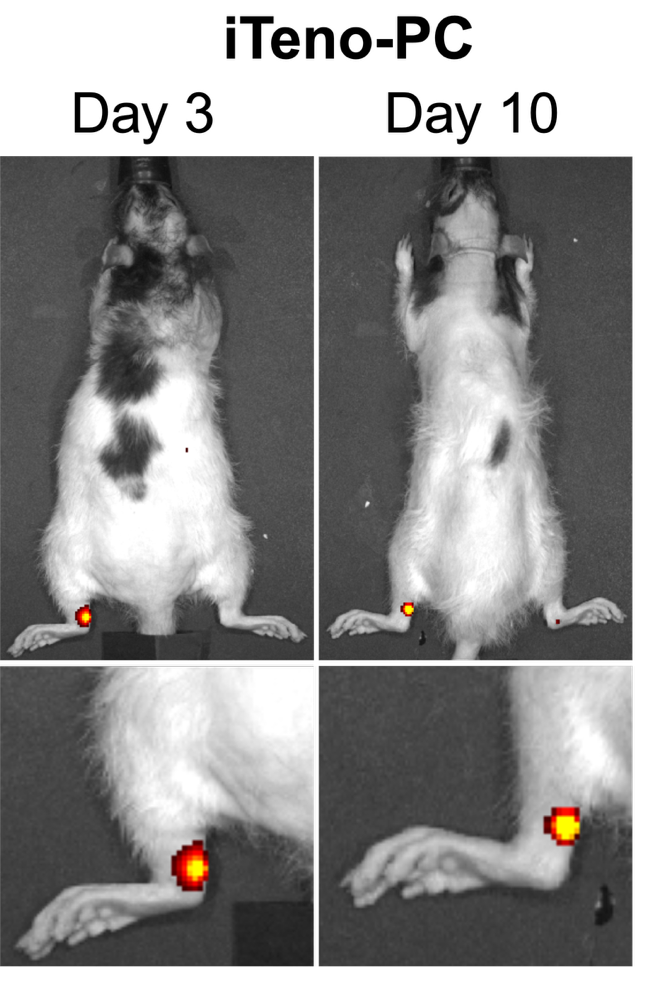

**Figure S5.** Representative IVIS images of iTeno-PC at days 3 and 10.
